# α-Synuclein driven cell susceptibility in Parkinson’s disease

**DOI:** 10.1101/2025.08.19.670819

**Authors:** Jonathan C. Breiter, Joseph S. Beckwith, Emma E. Brock, Joanne Lachica, Christina E. Toomey, Bin Fu, Mina Ryten, Lucien E. Weiss, Nicholas W. Wood, Sonia Gandhi, Michele Vendruscolo, Steven F. Lee

**Author notes:** Contributed equally to this work (Jonathan C. Breiter, Joseph S. Beckwith).

## Abstract

Early cellular events in Parkinson’s disease (PD) remain elusive. While aggregation of α-synuclein (αSyn) into Lewy bodies marks advanced pathology, smaller αSyn oligomers have been implicated in prodromal stages. Here we map αSyn oligomers at single-particle resolution in post-mortem brain tissue from Braak stage 3/4 PD cases and matched controls. Quantitative imaging of 9,882 neurons across four regions captured over 112 million αSyn oligomers. Mean intracellular α-Syn burden was unchanged between groups, but PD samples contained a higher fraction of neurons whose oligomer load exceeded a specific aggregation threshold. We term these aggregation-susceptible cells (ASCs). ASC enrichment in vulnerable regions supports a population-level model in which early pathology arises from a stochastic shift in cellular composition rather than altered αSyn aggregation kinetics. This human-tissue, large-scale dataset provides a quantitative framework for detecting ASCs and for testing population-level interventions in PD and related proteinopathies.

## Main Text

Lewy bodies (LBs)—intraneuronal inclusions rich in aggregated αSyn—are a defining pathological hallmark of PD, yet their role in disease initiation remains uncertain.^1–7^ While clinical severity correlates poorly with LB burden, evidence points to small soluble aggregates of αSyn as potential neurotoxic species in PD pathogenesis.^1,8–10^ However, despite decades of study, the prevalence of α-synuclein oligomers in the human brain, and the identity and distribution of the neurons most affected by them, have remained largely undefined.^9,11–13^

Biochemical studies first detected elevated soluble αSyn oligomers in PD patient biofluids and in dementia with Lewy bodies (DLB) cortex more than fifteen years ago.^9,14^ More recently, proximity ligation imaging visualised diffuse oligomeric pathology even in brain regions devoid of LBs and patient skin samples.^15–17^ However, a critical gap in knowledge remains: are αSyn oligomers a diffuse hallmark of prodromal PD, or do they emerge in a rare and expanding subset of vulnerable neurons? There is thus a need for quantitative surveys of αSyn aggregates across large numbers of individual neurons in early-stage human tissue.

Here we address this problem by applying a high throughput, single-aggregate microscopy method^18,19^ to post-mortem cortical, limbic and striatal tissue from individuals at Braak stages 3/4. By profiling more than 100 million phosphorylated αSyn puncta at subcellular Nyquist resolution, we identify a distinct subpopulation of neurons marked by elevated αSyn oligomer burden. We term these Aggregation Susceptible Cells (ASCs). This ASC subpopulation more than doubles in frequency in the acutely affected parahippocampal cortex but declines in the late stage affected caudate. Critically, we observe no change in aggregate size (intensity) distributions (**Figure S1**), suggesting that the key pathological transition is not a shift in αSyn assembly mechanisms but a probabilistic redistribution in the population of neurons that cross the threshold for pathological aggregation.

These findings refine current views of where and how early PD emerges. Rather than indicating a uniform per-cell change in αSyn accumulation or a distinct oligomer species, our data are consistent with an increased prevalence of ASCs as neurons whose αSyn oligomer burden exceeds an empirically defined aggregation-relevant threshold. By characterizing ASCs, we provide a cellular framework for localising early aggregation-prone subpopulations in human tissue. This work emphasises rare, stochastic cellular events that precede overt inclusions and offers quantitative readouts that may aid tracking of early pathology and, in future studies, inform intervention strategies.

### Transition from homeostasis to early disease state

LB presence affects only a very small subset of cells. Conservative estimates based on recent studies suggest that fewer than 0.1% of neurons in the *substantia nigra* develop LBs, with even lower proportions likely in cortical regions.^16,20–22^ Therefore, the thermodynamically driven formation of an LB in any one cell across the brain is an exceedingly rare event accounting for just 1 in 100 million, or fewer, cells (**Section S1**) (**Fig. 1A**). Therefore, remarkably few neurons contain intracellular aggregates of αSyn matching the classic definition of an LB.^23,24^ However, given that all cells produce αSyn, which multimerises readily as part of regular functioning in the healthy brain,^25–30^ they will likely all contain a range of smaller assemblies. Conceptually, each neuron can be considered a reaction vessel in which αSyn exists across a continuum of assembly states (**Fig. 1A)**. Under physiological conditions, αSyn is maintained in a dynamically regulated equilibrium of monomeric protein and largely soluble, non-pathogenic multimers.^27,28,31^ For LB formation to occur, this steady state must shift toward the accumulation of higher-order multimers that are aggregation competent^32–36^ **(Fig. 1A)**. In each region of the healthy brain, all neurons can be considered to exist on a continuum of intracellular αSyn oligomer concentration (**Fig 1B**), where most cells contain a homeostatically regulated amount of αSyn which makes the probability of each individual cell of forming a pathological aggregate minimal. Exceedingly few neurons contain a large amount of physiological αSyn, increasing the probability of the onset of runaway αSyn aggregation.^33–38^ We therefore define these cells as ASCs.

**Fig. 1.**
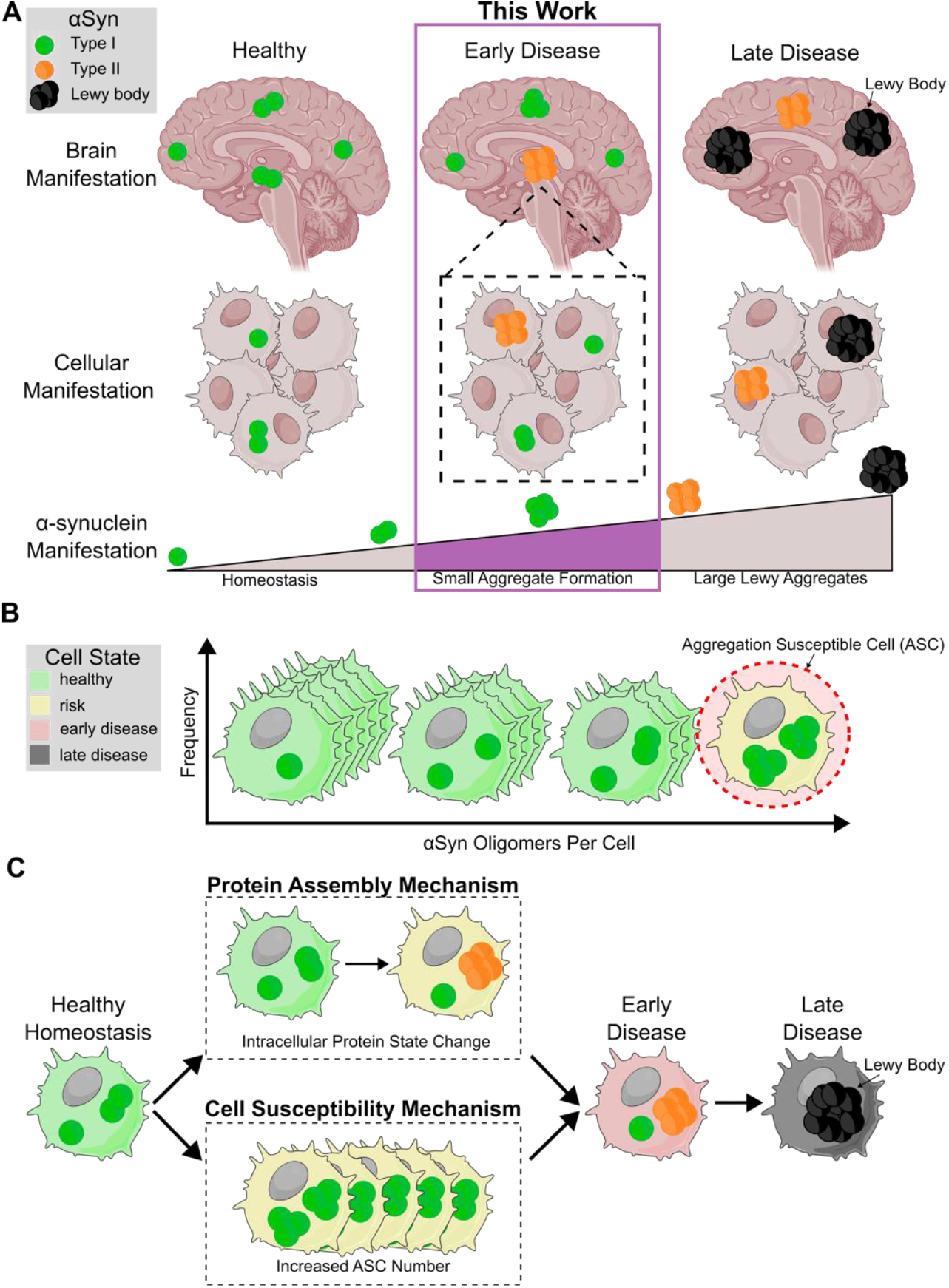
Conceptual framework for α-synuclein aggregation states and cell-population susceptibility in Parkinson’s disease (PD). **A**. PD progression at the brain, cell and αSyn level. Neurons in healthy brains contain a distribution of small αSyn aggregates which are homeostatically controlled. Early PD is defined an increase in aggregation propensity of a subset of aggregates (Type II). In late PD, these aggregates have matured into LBs. **B**. At the cell population level, cells with high aggregate concentrations are rare, while healthy cells with lower aggregate concentration are more frequent. The cells that have high aggregate concentrations are most susceptible to runaway aggregation, and we herein refer to them as Aggregation Susceptible Cells (ASCs). **C**. The initial transition from healthy cellular homeostasis towards disease can occur *via* two mechanisms. In the protein state change mechanism, there is an alteration of protein assembly kinetics, which permit pathological aggregate formation in any cell within the population, not exclusively in ASCs. By contrast the cell aggregation susceptibility mechanism involves an increase in the proportion of ASCs at the population level, thereby elevating the likelihood of pathological aggregate formation specifically within this subpopulation.

Two mechanisms can account for the early transition of individual neurons from physiological αSyn homeostasis to pathological aggregation **(Fig. 1C)**. In the first mechanism, which impacts protein-state change, αSyn undergoes a transformation that renders it intrinsically more prone to aggregation. Here, the distribution of the population of cells remains unchanged, but the aggregation propensity within individual neurons increases. This would result in either aggregates with altered structural or biochemical properties appearing in disease tissue relative to controls, or in uniquely high αSyn-containing cells.^19^ Such a mechanism is supported by evidence that point mutations, splice variants, or post translational modifications can shift αSyn toward more aggregation prone states, thereby highlighting protein-state change as a sufficient mechanism for PD onset.^39–43^ In the second mechanism of increased cell aggregation susceptibility, the aggregation pathway remains biophysically unaltered, but a shift occurs at the level of the cell population. In this scenario, a greater fraction of neurons begins to accumulate physiological αSyn to levels that surpass a critical threshold, making them susceptible to spontaneous primary and runaway secondary nucleation events. This reflects a probabilistic population level transition in which stochastic fluctuations in αSyn concentration tip more cells into an aggregation prone state, even without any intrinsic change to the protein itself.

To describe this phenomenon, ASCs are neurons whose αSyn content places them near or beyond the threshold required for spontaneous aggregation. In a healthy brain region, neurons can be viewed as a heterogeneous population, each containing a different amount of soluble αSyn (**Fig. 1B**). The majority maintain homeostatic levels, remaining below the critical threshold and therefore at minimal risk for aggregation. A small minority, however, may accumulate higher concentrations due to transcriptional, translational, or clearance related fluctuations.^44^ These rare cells, ASCs, are the most vulnerable to crossing into pathological states, particularly under conditions that subtly shift population level distributions.

A direct link between intracellular αSyn oligomer burden and neuronal susceptibility to aggregation remains the critical missing link in early PD pathogenesis. Despite clear conceptual models, this connection has been difficult to test in human tissue, largely due to the technical challenge of detecting small aggregates at scale. *In vitro* studies have demonstrated both molecular and population level mechanisms under controlled conditions^42,45–48^ but this question has not been fully resolved *in situ*.

## Results & Discussion

### Measurement of intracellular αSyn concentration across the brain

In order to study the mechanism of initial PD manifestation, [αSyn Aggregate]_Cell_, herein defined as aggregate concentration, was quantified in fourteen early stage (Braak ¾) PD patients and thirteen demographics-matched healthy controls (HCs) across three brain regions. We chose to quantify the Serine 129 phosphorylated form of αSyn as this post-translational modification (PTM) shows a correlation to changes in aggregation kinetics,^49,50^ is heavily associated with disease,^49,51–53^ is used commonly as a neuropathological marker for all synucleinopathies,^54,55^ and has previously shown strong compatibility with our ASA-PD protocol (SI **Figure S2, Section S3**).^19^ The three brain regions mapped can be ordered in terms of disease severity according to Braak^56,57^: the Frontal Cortex (unaffected in PD), the Parahippocampal Cortex (mildly affected) and the Caudate Nucleus (moderately affected) (**Fig. 2A + Section S2**). Quantitative bioimaging was conducted according to our ASA-PD protocol^,19^ which yields the 3D location of single aggregates in relation to MAP-2 positive neurons and aggregate brightness, which can be used as a proxy for size^19^ (**Figure 2 B, Figure S7**). We observed a total of 4,327 neurons (2,407 HC, 1,920 PD) in the Frontal Cortex, 3,136 neurons (1,540 HC, 1,596 PD) in the Parahippocampal Cortex and 2,419 neurons (1,229 HC, 1,190 PD) in the Caudate Nucleus containing a total of 112,097,781 pS129 αSyn oligomers (**Figure 2C, Table S1**). From this data, per-cell aggregate concentration was computed as the number of diffraction-limited pS129 αSyn puncta inside neurons, normalised to the volume of the neuron (**Figure 2C**) but not local density, as it did not vary across patients or disease state (**Figure S3**). Aggregation Susceptible Cells, ASCs, were defined based on an aggregate concentration threshold. This threshold is iteratively scanned across the concentration distributions for quantitative observations of changes (**Section S5**). We determine the proportion of ASCs *versus* all observed cells for each brain region (**Figure 2C**).

**Fig. 2.**
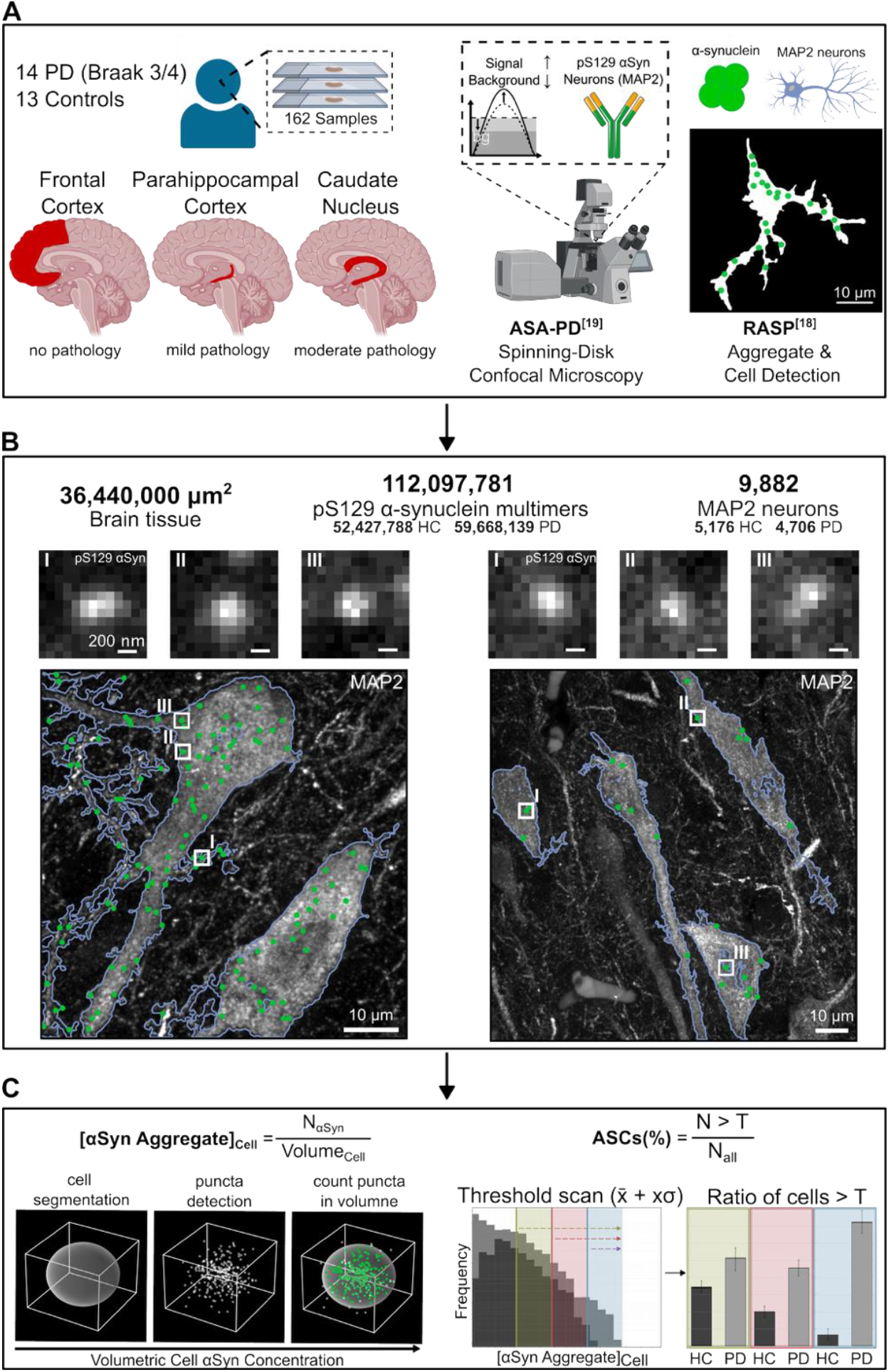
High-throughput quantitative bioimaging detected 112 million αSyn oligomers within control and PD neurons. **A**. Samples from three brain regions with a pathological severity gradient. Samples were prepared following the ASA-PD procedure^19^ stained with an anti-pS129 αSyn antibody (AB_2270761), and imaged using a bespoke high-NA spinning-disk confocal microscope. Subsequent high-throughput quantitative single-aggregate microscopy quantified the 3D colocalisation of αSyn assemblies with the neuronal marker MAP2 (AB_2936822).^18,19^ **B**. Representative images of MAP-2 positive neurons in the cerebral grey matter. Green puncta show localisations of diffraction-limited pS129 αSyn assemblies, which are highlighted in the crop-ins. **C**. Aggregate concentration was calculated by dividing the number of observed diffraction-limited αSyn assemblies per cell over the volume of the cell. The percentage of ASCs (%), is calculated by applying a scanning threshold (x = 0– 3 in steps of 0.05) to the distributions of aggregate concentration and taking a ratio of cells > threshold for both HC and PD distributions at each iteration (see Section S5).

### Intraneuronal αSyn concentration is not increased in early PD

The distributions of aggregate concentration in the three brain regions in increasing order of disease severity after Braak^56,57^ is shown in **Figure 3A**. These distributions do not significantly differ between the healthy control (HC) and PD samples for neurons. The right-most plot in **Figure 3A** quantifies a ratio of PD over HC aggregate concentration and shows no clear differences between PD and HC in any brain region examined here, with the ratio in all brain regions equalling 1, within error (see **Sections S4 + S6**). This data shows that, irrespective of a brain region’s Braak stage disease severity, the intraneuronal concentration of small, soluble assemblies of αSyn is not significantly increased in PD compared to a HC group. These findings oppose the protein state change mechanism (**Figure 1D**), as there is no significant subset of cells with uniquely high concentrations, as would be expected if protein assembly kinetics changed fundamentally within a subset of cells. Additionally, a significant change in aggregate brightness would be predicted in heavily affected regions, as demonstrated before in late Braak stage PD.^19^ Importantly, the change in the state of individual αSyn aggregates is entirely absent in our data (**Figure S1**). These findings argue against the protein-state change model operating in early disease (Braak ¾), indicating that altered protein assembly kinetics do not underlie the initial pathological transition.

**Fig. 3.**
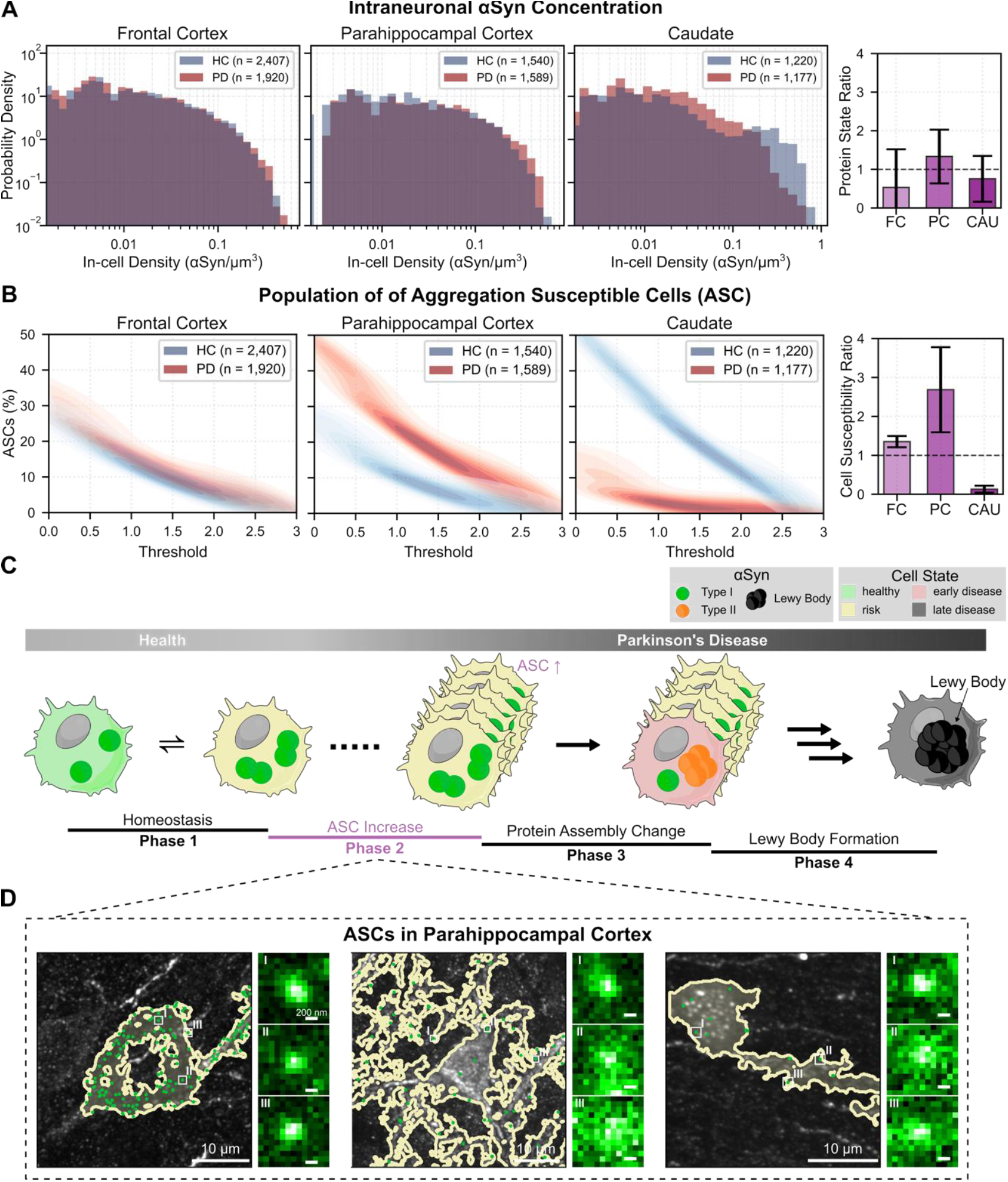
ASC prevalence is increased in parahippocampal cortex, unchanged in frontal cortex, and decreased in caudate in PD vs controls. **A**. Probability density distributions of the aggregate concentration across the different brain regions imaged. The summary plot shows the Mean ± SD ratio of PD/HC aggregate concentration. **B**. The sub-population of ASCs as a function of a changing threshold on aggregate concentration where increasing threshold corresponds to more extreme values within the aggregate concentration distribution (see **Fig. 2C, Section S6**). The summary plot shows the Mean ± SD ratio PD/HC of ASCs (see **Section S4**). **C**. Mechanistic representation of the observed mechanism of early-PD manifestation. An increase in the fraction of ASCs is observed prior to a change in protein state. After the increase in ASC abundance, αSyn aggregates will undergo a transition to seed-competent aggregates (state-change) in a subset of ASCs, which form Lewy Bodies in late-stage PD. **D**. ASCs and the aggregates contained within them across the parahippocampal cortex of three PD patients. Crop-ins highlight the diffraction-limited aggregates of pS129 αSyn.

### The fraction of neurons with high intraneuronal αSyn concentration is increased in early PD

Quantifying the percentage of ASCs for a given brain region is done by setting a threshold (T) on intracellular αSyn aggregate concentration (**Section S5**). The probability density of ASCs (%) as a function of T is shown in **Figure 3B**. In the frontal cortex, the proportion of ASCs in HCs and PD patients are largely the same across T, meaning no increase in the relative proportion of ASCs is associated with Braak stage ¾ PD in the frontal cortex. In the parahippocampal cortex, a clear separation between the proportion of ASCs in PD compared to HC can be observed across the majority of thresholds (T). Up until a threshold of x̄ + 1.5σ, the average proportion of ASCs in early PD patients’ parahippocampal cortex is ∼2.5 fold higher than in HCs. Thus, in a region where disease is currently advancing, an increase in ASCs is consistent with an increased likelihood of LB formation, as more cells have a high probability of further aggregation. In the caudate, where PD has already progressed to moderate severity, including neuronal death and LB formation,^6,56,57^ there is a depletion of the proportion of ASCs in PD patients compared to HCs. We propose this is due to high aggregate concentration neurons already having formed LBs, depleting the measurable aggregates by sequestering them into LBs, as well as depleting the population of ASCs. This is also evidenced by an overall decrease in αSyn aggregate density in this brain region in PD compared to HC (**Figure S3**). Similar findings were also observed in microglia taken form the same areas and cases (**Figure S4**). Overall, these observations are in agreement with previous work suggesting that LB formation may be a protective mechanism, insofar as LBs sequester soluble αSyn assemblies, depleting their global concentration.^1,2,9,16,58^. We confirmed that a sufficient number of cells were sampled to ensure statistical robustness (**Figure S5**), and measurements were consistent across unaffected brain regions (**Figure S6**).

These data are supportive of the cell aggregation susceptibility mechanism (**Figure 1C**). This suggests that an increase in the number of ASCs is the first manifestation of αSyn-related pathology in the early stages of PD. This may mark an early shift toward a disease-permissive state, which is predominantly characterised by the intracellular assembly of LBs in a subset of the ASC neurons (**Figure 3C**). Thus, the initial increase in the number of ASCs precedes the protein change that generates seed-competent aggregates that then can grow into LBs. We highlight parahippocampal ASCs from three mid-stage PD patients to showcase the morphological heterogeneity of neurons which are ASCs and show the variable subcellular αSyn aggregate distribution within ASCs (**Figure 3D**).

Our data highlight an early, cell population level shift in cellular aggregation susceptibility in Braak stage ¾ brain tissue. It is important to note that all tissue examined here precedes the classical molecular pathology of widespread LB presence observed in Braak stages 5/6. It is during these later stages that a conformational or kinetic change in the αSyn protein is likely to become more dominant, proceeding via the proposed protein-state change mechanism. Such an interpretation is substantiated by recent work in late-stage PD using our ASA-PD protoco^l19^ as well as extensive previous work *in vitro*, in cell and animal models, and in patient samples.^9,10,13,17,18,29–31,34,45,47,48,51,58,59^

Our findings here characterise ASCs as key contributors to the early stages of PD. The increased prevalence of ASCs we observe in earlier disease stages represents a permissive cellular landscape upon which such protein-level changes later act. These findings suggest that increased cell aggregation susceptibility may precede detectable changes in αSyn aggregation kinetics. Kinetic studies of αSyn aggregation and genetic variants of disease, such as PTMs or the A53T mutation, increase aggregation propensity and likely directly engage protein-state change^,38–43,47^ effectively bypassing the initial cell-population change. Under this newly proposed framework, many animal models of synucleinopathy will similarly bypass the here observed early shift in cell aggregation susceptibility, which is evident in the absence of protein-state change, progressing straight to a disease-permissive state through engineered alterations of αSyn assembly. While these models remain valuable for investigating disease mechanisms downstream of runaway αSyn aggregation, they may fail to capture key early events in idiopathic PD.

The interpretations of the evidence presented here have limitations. Firstly, our focus was exclusively on αSyn phosphorylated at Serine 129. Although this is a widely accepted disease marker,^2,38–43,47,51–53^ it does not reflect the full spectrum of αSyn aggregate species relevant to early pathogenesis. Second, although this is a large-scale study, imaging over 36,440,000 µm^2^ with Nyquist sampling, the number of brain regions (three) and post-mortem cases (14 PD, 13 HC) remains limited. Our sampling thus covers only a subset of the anatomically, temporally and individually diverse landscape of the disease. Third, we examined only sporadic PD. Familial forms, particularly those driven by strong genetic components, may be more likely to progress via the protein-state change mechanism, reflecting intrinsic alterations in αSyn assembly rather than shifts in cellular aggregation susceptibility shown here. Future studies should aim to complete the picture by investigating the presence and abundance of ASCs in these genetic forms of PD.

Nonetheless, our findings suggest that changes in ASC prevalence precede detectable alterations in αSyn aggregation kinetics. This supports a framework in which cell-level aggregation susceptibility emerges before molecular transformation, providing a basis for future studies that integrate these two axes of disease onset and progression.

## Author contributions

Conceptualization: SFL, MV, SG, NWW, MR, JCB, JSB

Methodology: EB, JCB, JSB, CET, JL, BF

Investigation: EB, JCB, CET, JL

Visualization: JCB, JSB, BF, EB

Data Analysis: JSB, JCB, SFL, BF, EB, LW

Funding acquisition: SFL, MV, SG, NWW, MR

Supervision: SFL

Writing – original draft: SFL, JSB, JCB

Writing – review & editing: SFL, JSB, JCB, MV

## Data availability

Data supporting this study is available at three levels of complexity.

1. **Raw imaging data** is archived independently at the *Imaging Data Resource (IDR)*, with a DOI currently being generated (∼2 TB). [DOI under review at IDR].
2. **Processed data** quantifying the cellular density of αSyn is available as a structured database on Zenodo (https://doi.org/10.5281/zenodo.16421701)
3. **Exemplar data and analysis code**, suitable for reproducing key findings, are provided alongside this publication “pyRASP_copy_for_paper.zip” and on Zenodo at https://doi.org/10.5281/zenodo.16421701.

## Code availability

Code supporting this study is available both at GitHub (https://github.com/TheLeeLab/pyRASP) and an archived version with example data is available at Zenodo at https://doi.org/10.5281/zenodo.16421701.

## Supplementary Information

Figs. S1 to S9

Sections S1 to S7

Table S1

## Materials & Methods

### Post-mortem sample selection

All brain samples used in this study were provided by the Multiple Sclerosis and Parkinson’s Brain Bank, Imperial College London. Ethical approval and informed consent were provided for all cases. Selection of cases and controls was performed based upon neuropathological diagnosis of each case (Control or Parkinson’s disease) after undergoing detailed diagnostic screening by a qualified neuropathologist. Cases were chosen to have no dementia incidence, mapped to the Brainstem predominant or limbic transitional stage of McKeith staging criteria, post-mortem delay (less than 24 hours) and where possible an absence of any confounding pathology. The exception being that some HC had minimal age-related changes. The two cohorts, namely HC and PD, had comparable mean age of death (age_HC_ = 83.4; age_PD_ = 79.2) and both groups had a higher representation of male subjects relative to female (ratio_HC_ = 2.6; ratio_PD_ = 3.5). All case demographics can be found in more detail in Supplementary **Table S1**.

A key distinction of this study is its focus on Braak stages 3/4 in regions that are only mildly affected, allowing us to investigate the mechanisms driving the earliest phases of disease progression. By contrast, much previous work has centred on late-stage disease (Braak 5/6), where the initial changes have likely already occurred.

### Sample Preparation - FFPE Human Brain Slices

Formalin-fixed paraffin-embedded (FFPE) tissue sections were obtained from the cingulate cortex (see tables S2 and S3) and cut to 8 μm thickness. FFPE sections were baked at 37°C for 24 hours followed by 60°C overnight. Sections were deparaffinized in xylene and rehydrated using graded alcohols. Non-specific binding was blocked with 10% bovine serum albumin (BSA) solution in PBS for 30 minutes. Heat mediated epitope retrieval was undertaken with tissue sections pressure cooked in citrate buffer at pH 6 for 10 minutes. Tissue sections were incubated with primary antibodies; anti-phosphorylated alpha-synuclein (ab59264, AB_2270761, Abcam, 1:200); and either ionized calcium-binding adapter molecule 1 (GT10312, Thermo Fisher, AB_2735228, 1:500) or Microtubule-Associated Protein 2 (ab254143, Abcam, AB_2936822, 1:500) for 1 hour at room temperature. The sections were then washed three times for five minutes in PBS followed by the corresponding AlexaFluor secondary antibodies (anti-rabbit 568, A11011, AB_143157, Thermo Fisher, anti-mouse 488, A11001, AB_2534069, Thermo Fisher, all at 1:200) for an additional hour at room temperature in the dark. Sections were then washed three times for 5 minutes again in PBS and incubated in Sudan Black B (0.1% for 10 minutes, 199664-25G, Sigma Aldrich). Removal of Sudan Black B occurred with three washes in 30% ethanol (E7148-500ML, Sigma Aldrich) before being mounted with Vectashield+ (Vector Labs, H-1900, VWR, 50 mm × 24 mm #1 thickness, Catalogue Number 48404-453) for imaging. Sections were stored at 4°C until imaging was completed.

### Optical Setup

The microscope used to image the FFPE human brain slices was a spinning disk confocal microscope (3i intelligent imaging). The microscope was equipped with a 200 mW, 488 nm laser (LuxX) and a 150 mW, 561 nm laser (OBIS). These lasers were housed in a beam combiner (3i intelligent imaging), which focused them into an optical fiber which sent the illumination light into a field flattener (Yokogawa-Uniformizer for CSUW). The excitation light was then passed into a spinning disk unit (50 μm sized pinholes, Yokogawa CSU-W1 T2 Single-Molecule Spinning Disk Confocal, SoRa Dual Microlens Disk) and then the microscope body (Zeiss Axio Observer 7 Basic Marianas™ Microscope with Definite Focus 3) using a dichroic mirror (FF01-440/521/607/700, Semrock). The fluorescence was filtered using either a FF01-525/45-25-STR filter (Semrock) in the case of 488 nm excitation or a FF02-617/73-25-STR filter (Semrock) in the case of 561 nm excitation. The fluorescence was then focused onto one of two sCMOS cameras (Prime 95B, Teledyne Photometrics). The objective lens was a Zeiss oil immersion objective (Alpha Plan-Apochromat 100x/1.46 NA Oil TIRF Objective, M27), and the microscope was controlled using a PC (Dell-Acquisition Workstation 310R) and software (Slidebook, 3i intelligent imaging). The 561 nm power density was 260 ± 12 W/cm^2^, and the 488 nm power density was 250 ± 100 W/cm^2^. The 488 nm power density (cell label channel) varied more as this was tuned per day to avoid saturating the camera in any pixel, depending on label density.

### Brain imaging data acquisition

Brain imaging was conducted according to previously established protocol.^1^ Briefly, three random locations were chosen for each brain tissue section in the grey matter. At each location, a 3 × 3 grid of 132 μm (x) × 132 μm (y) × 12 μm (z) z-stacks were acquired with 100 ms exposure time. This amounts to a total of 675 images per patient per cell type per brain region with an area of 5,645,376 μm^2^ covered on each sample.

### Camera gain calibration

To convert the pixel value to photons in a sCMOS camera, we recorded a series of image sequences at 7 different intensity levels (20,000 frames per intensity level) with uniform illumination, including one level at no illumination for the calculation of camera offset. For every pixel, the mean and variance were calculated across the 1,000 frames, generating 7 different variance and mean values corresponding to the 7 non-zero illumination intensities. The camera offset per pixel was determined as the mean pixel value in the non-illuminated frame. The camera gain per pixel, expressed in photoelectrons per count, was determined by calculating the slope between the 7 variance and mean values per pixel, and subtracting the non-illuminated frame offset.^2^ Software available for this purpose is available at https://doi.org/10.5281/zenodo.10475643.

### Brain imaging data analysis

Aggregate detection proceeded using the RASP pipeline described in Fu *et al*.^3^ To briefly summarise, images underwent a high-pass kernel, obtained through the difference between the original image and a Gaussian-blurred image (σ = 1.4px), followed by a Laplacian-of-Gaussian (LoG) kernel (σ = 2px). Puncta were selected as pixels in the top 95th percentile of brightness, and these were then accepted or rejected based on their integrated gradients and flatness.^3^ These gradient and flatness thresholds are determined based on negative control data for each round of imaging.

For cell mask detection, each 2D image was enhanced using a difference-of-Gaussian filter, with σ_1_ = 2px and σ_2_ = 60px. A binary mask of the image was created using a Yen threshold.^4^ This binary mask then had a binary opening operation applied to it with a disk morphology of radius 1 pixel, which was then followed by a binary closing operation with a disk morphology of radius 5 pixels. When all of the images in a volumetric stack had binary masks generated in this way, the scikit.image function binary_fill_holes^5^ was applied to the full 3D volume, after which small holes of <100 voxels were removed. Post this, 3D objects below a specified cell size were removed to end up with the final cell masks used for analysis.

