## Supplementary Information for "α-Synuclein driven cell susceptibility in Parkinson’s disease"

**Affiliations:**
[1]Yusuf Hamied Department of Chemistry, University of Cambridge, Lensfield Road, Cambridge, CB2 1EW, UK
[2]Centre for Misfolding Diseases, Yusuf Hamied Department of Chemistry, University of Cambridge, Cambridge CB2 1EW, UK
[3]Aligning Science Across Parkinson's (ASAP) Collaborative Research Network, Chevy Chase, MD 20815, USA
[4]The Queen Square Brain Bank for Neurological Disorders, Department of Clinical and Movement Neuroscience, UCL Queen Square Institute of Neurology, London WC1N 3BG, U.K.
[5]The Francis Crick Institute, 1 Midland Road, London, UK
[6]Department of Neurodegenerative Diseases, UCL Queen Square Institute of Neurology, London WC1N 3BG U.K.
[7]UK Dementia Research Institute at the University of Cambridge, Cambridge CB2 0AH, U.K.
[8]Department of Clinical Neurosciences, School of Clinical Medicine, University of Cambridge, Cambridge CB2 0SP, U.K.
[9]Department of Genetics, University of Cambridge, Cambridge CB2 3EH, UK.
[10]Department of Engineering Physics, Polytechnique Montréal, Montréal, Québec H3T 1J4, Canada
[11]Department of Clinical and Movement Neurosciences, UCL Queen Square Institute of Neurology, London WC1N 3BG, U.K.

†Contributed equally to this work

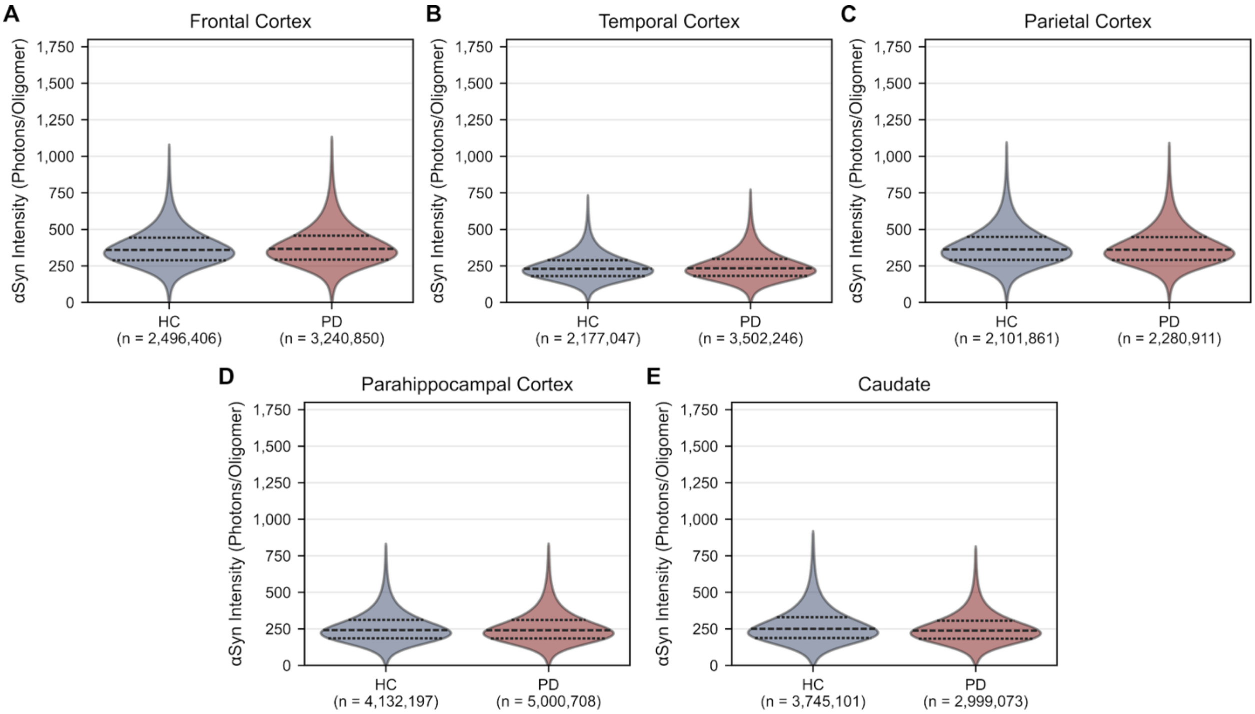


**Figure S1. There is no difference in aggregate intensity between PD and HC.** Intensity distributions of diffraction-limited αSyn aggregates detected in n = 13 HCs and n = 14 PD patients in **A.** frontal cortex **B.** temporal cortex **C.** parietal cortex **D.** parahippocampal cortex and **E.** caudate across two repeat imaging runs for each region each disease state. N-numbers represent the total number of aggregates detected across both repeats per region.

The dotted horizontal line in violins represents the median, with the upper and lower lines as the interquartile range (IQR).

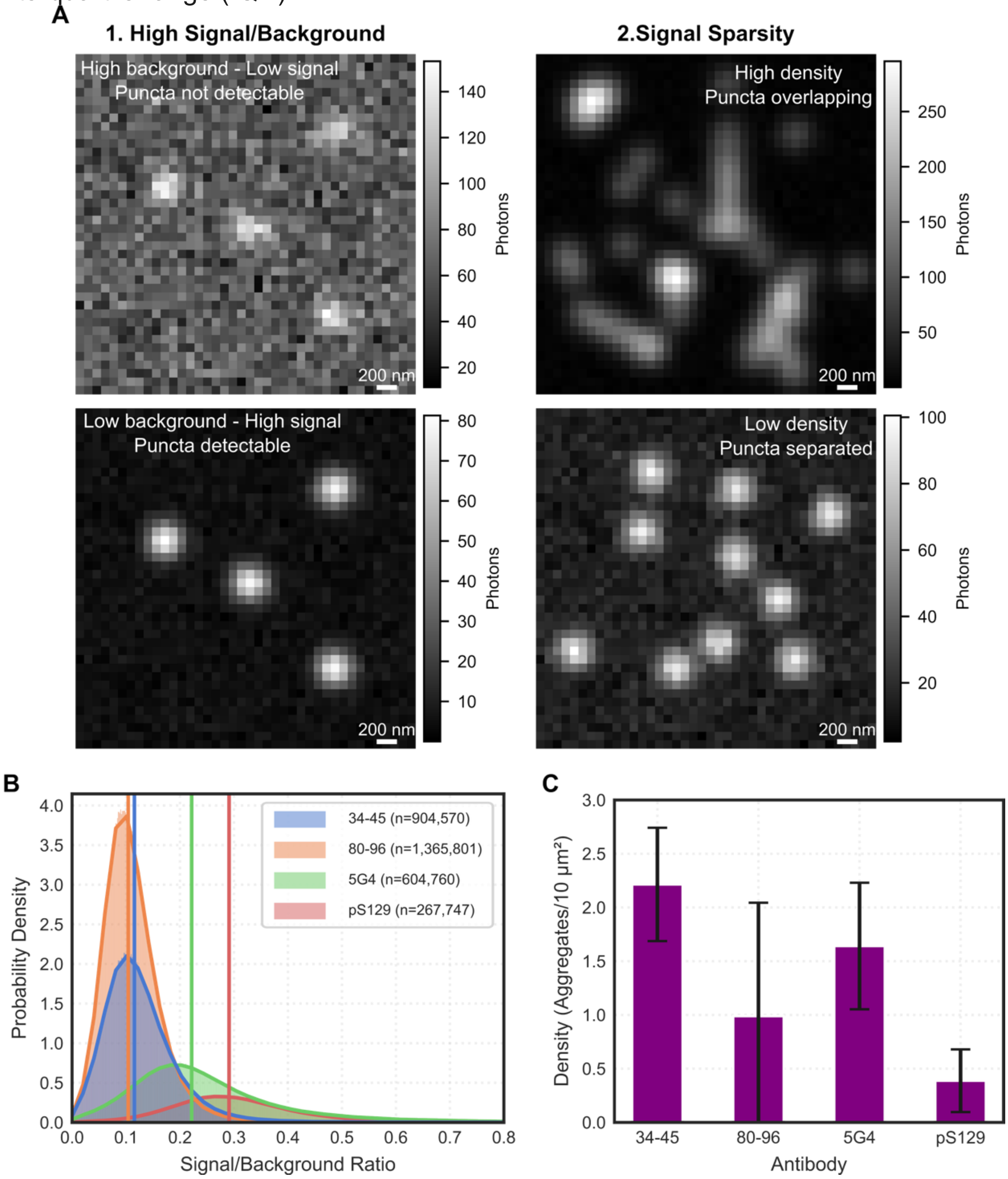


**Figure S2. Anti-pS129 antibody is most suitable for ASA-PD. A.** Simulated microscopy image-representation highlighting the two data quality requirements for ASA-PD and RASP.[1,2] Firstly, signal needs to be clearly separated from background intensity with a high signal/background ratio to detect individual diffraction-limited puncta. Secondly, signal needs to be sparse enough for spatial separation between point spread functions for accurate resolution of the location and quantity of puncta. **B.** Four αSyn antibodies tested in PD Braak 3/4 (n = 1) and HC (n = 1) brain. Signal over background ratio of oligomeric protein assemblies as detected by antibodies 34-45 (AB _2650701), 80-96 (AB_2650688), 5G4 (AB_2716647) and pS129 (AB_2270761). Vertical lines represent the median of each distribution. N-numbers represent the total number of aggregates detected by the antibody. **C.** Mean ± standard deviation density of aggregates produced by the four antibodies in the human brain. The pS129 antibody shows the highest signal/background ratio and the lowest density of diffraction-limited αSyn aggregates. Low background and high spatial separation of local maxima is essential for successful small aggregate detection.[1] Additional quantitative information on the requirements for the antibody election can be found in our previous work.[2]

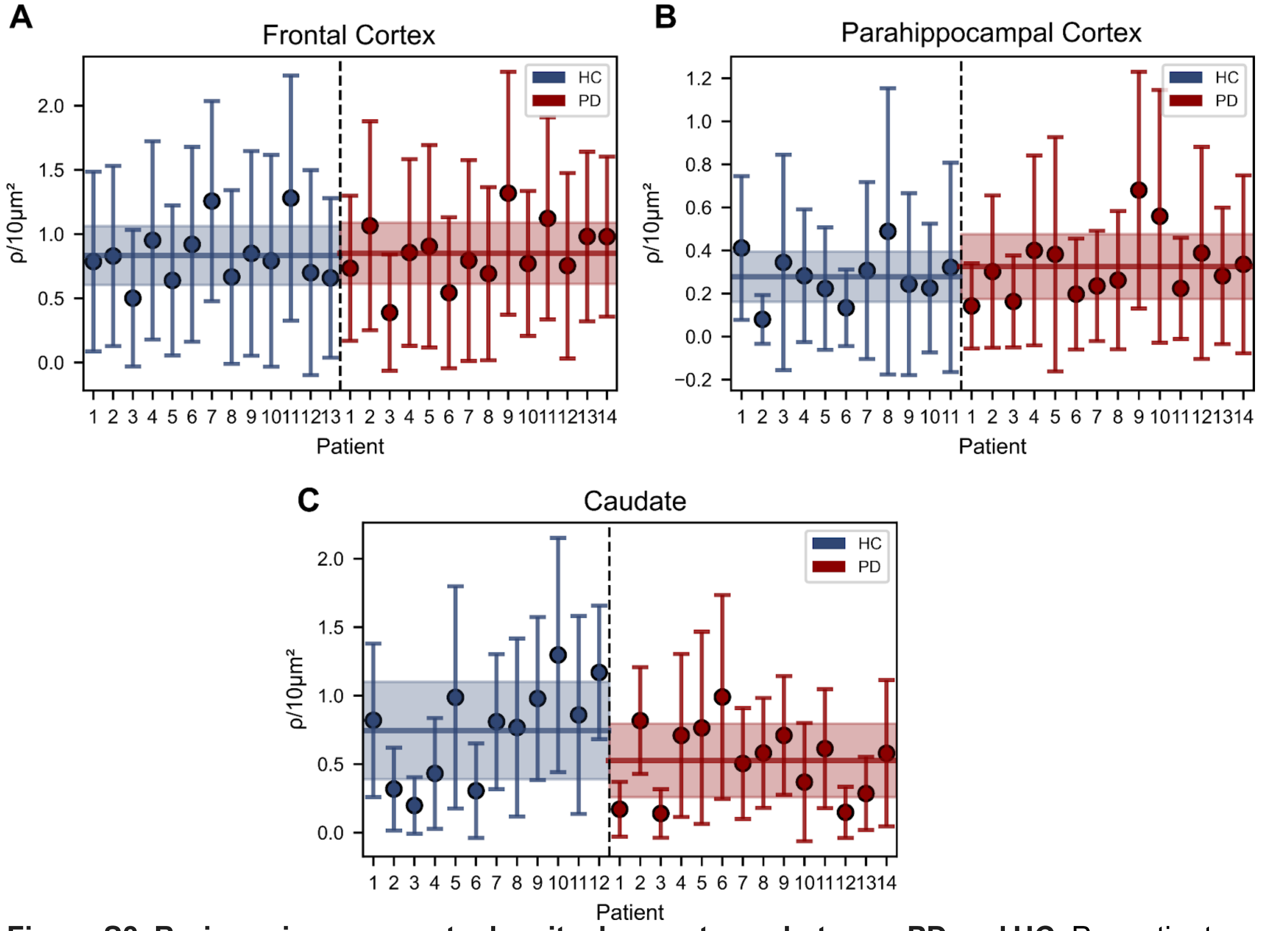


**Figure S3. Brain region aggregate density does not vary between PD and HC.** Per patient average density of αSyn oligomers per 10 μm² in the frontal cortex (**A.**), parahippocampal cortex (**B.**) and caudate (**C.**). Horizontal lines represent the mean ± SD across patient means. No significant difference is observed between individuals within the HC or PD group per region or between the HC and PD group averages in each region.

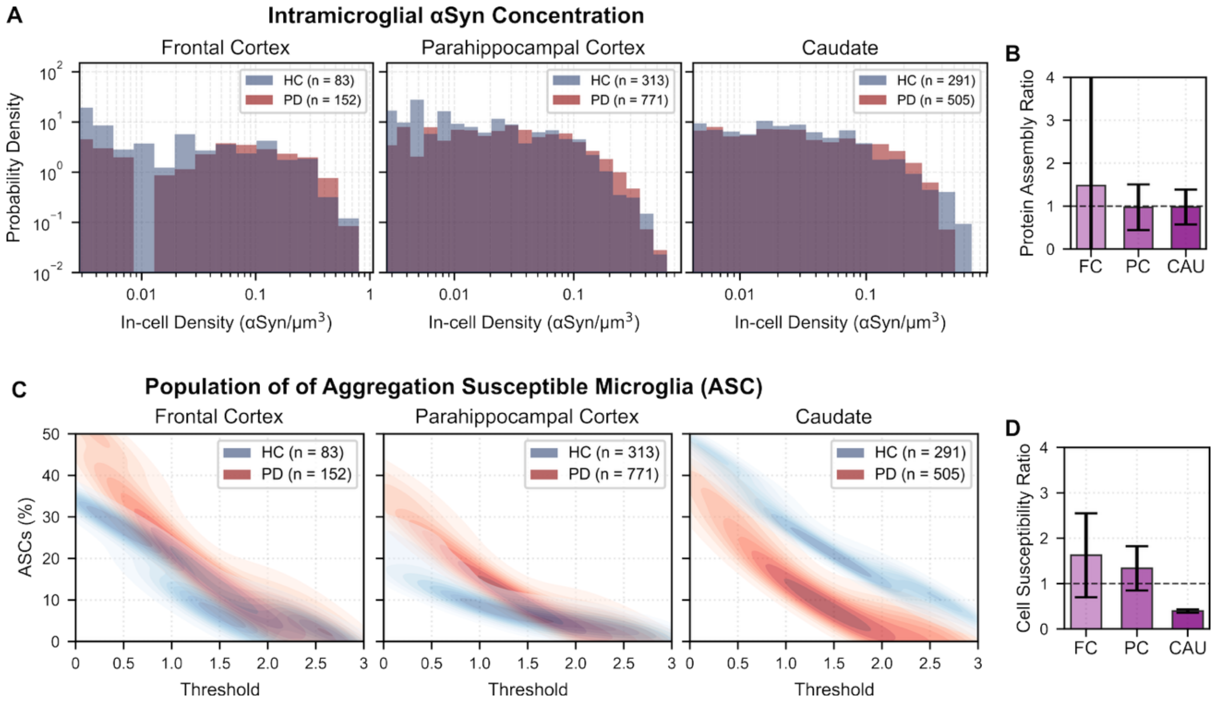

**Figure S4. Evidence for ASCs in microglia. A.** Probability density distributions of the aggregate concentration per microglia across three brain regions. **B.** The summary plot shows the Mean ± SD ratio of PD/HC aggregate concentration as the protein assembly ratio inside microglia. **C.** The sub-population of Aggregation Susceptible Cells (ASCs) of microglia as a function of a changing threshold on aggregate concentration, where increasing threshold corresponds to more extreme values within the aggregate concentration distribution (See **Figure 2 C**). **D.** The summary plot shows the Mean ± SD ratio PD/HC of ASCs of microglia as the cell susceptibility ratio.

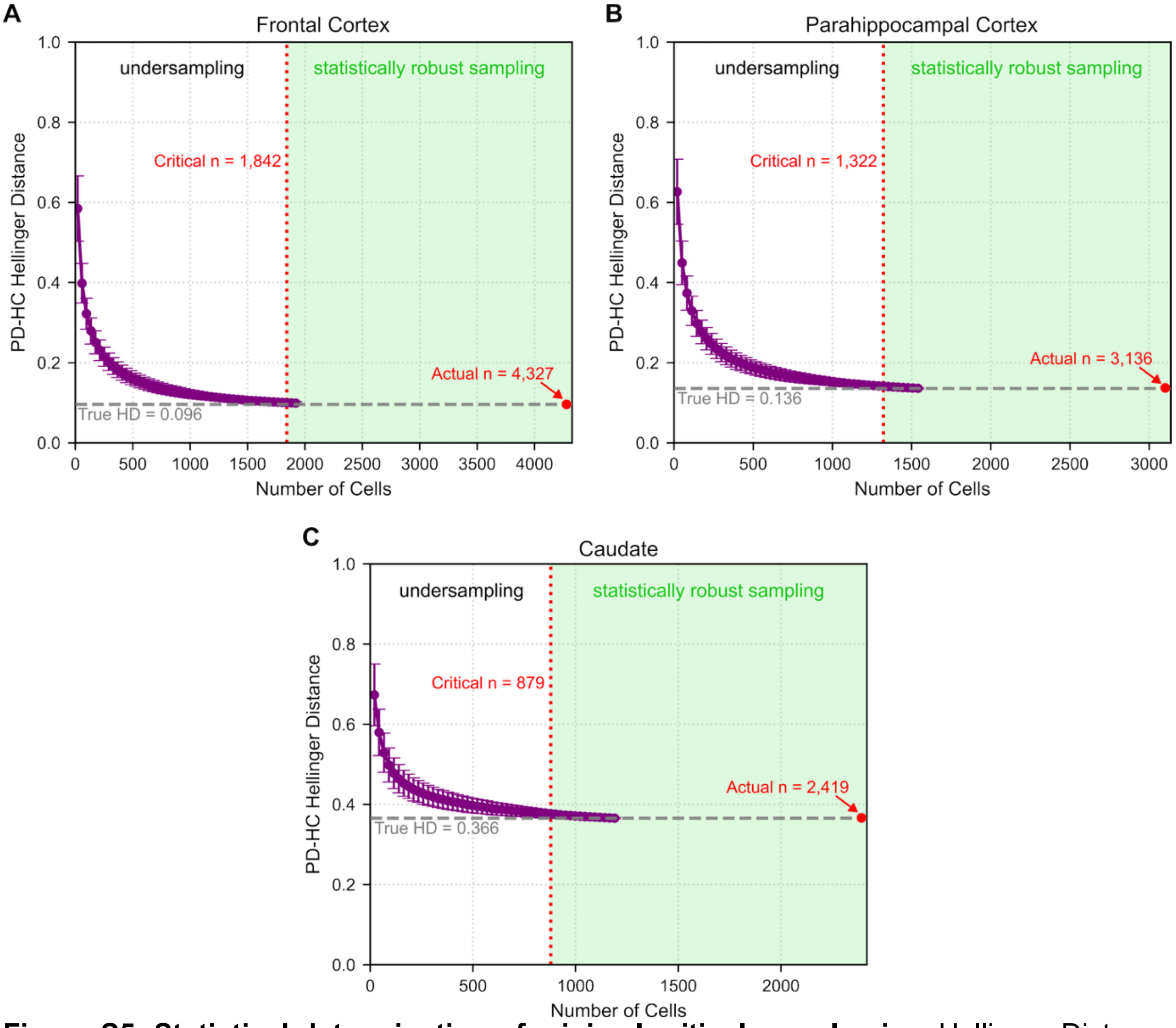


**Figure S5. Statistical determination of minimal critical sample size.** Hellinger Distance (HD) between PD and HC distributions in Figure 3 (**B.**) as a function of subsampled cells according to parametric HD[3,4] in frontal cortex (**A.**), parahippocampal cortex (**B.**) and caudate (**C.**). The minimum number of cells required to observe the true Hellinger distance between HC and PD is determined by subsampling the cell aggregate concentration distributions at decreasing n-numbers with 5,000 iterations per step and determining the initial intersection of mean ± SD across iterations of a subsampling step with the true observed HD. This shows the minimum number of neurons that need to be observed and quantified across PD and HC groups to power the observations made in **Figure 3 B**. Some points on true HD line are omitted for clarity to guide the eye.

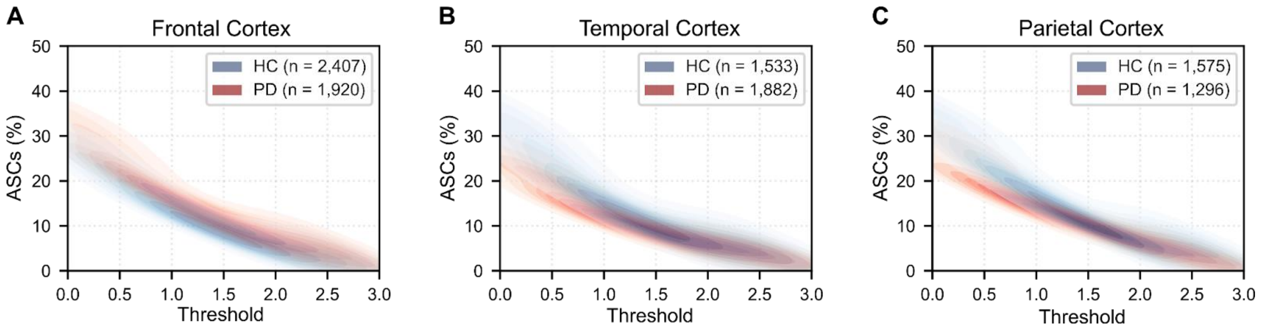


**Figure S6. There is no difference in ASC abundance in unaffected brain regions.** The percentage of Aggregation Susceptible Cells (ASCs) plotted against the scanning threshold set on the joint distribution of αSyn concentration per cell in the frontal (**A.**), temporal (**B.**) and parietal (**C.**) cortices in PD and HC samples. All three regions are considered to be unaffected by PD pathology in Braak stage 3/4.[5,6]

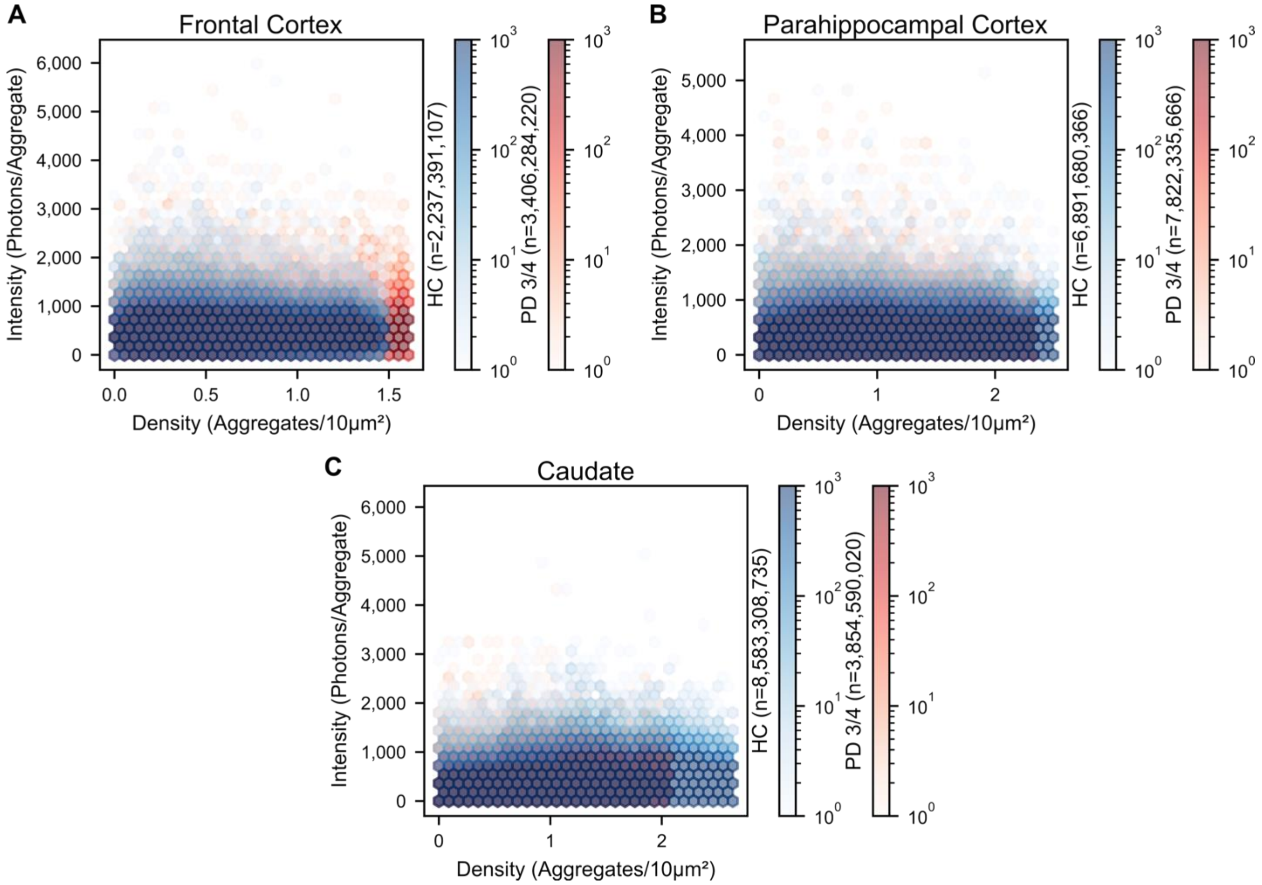


**Figure S7. Intensity and cellular density of aggregates show no correlation.** Correlation between the local density of diffraction-limited αSyn aggregates and the intensity per aggregate in PD and HC frontal cortex (**A.**), parahippocampal cortex (**B.**) and caudate (**C.**). The number of aggregates per bin is shown by the lookup colour bar next to each graph.

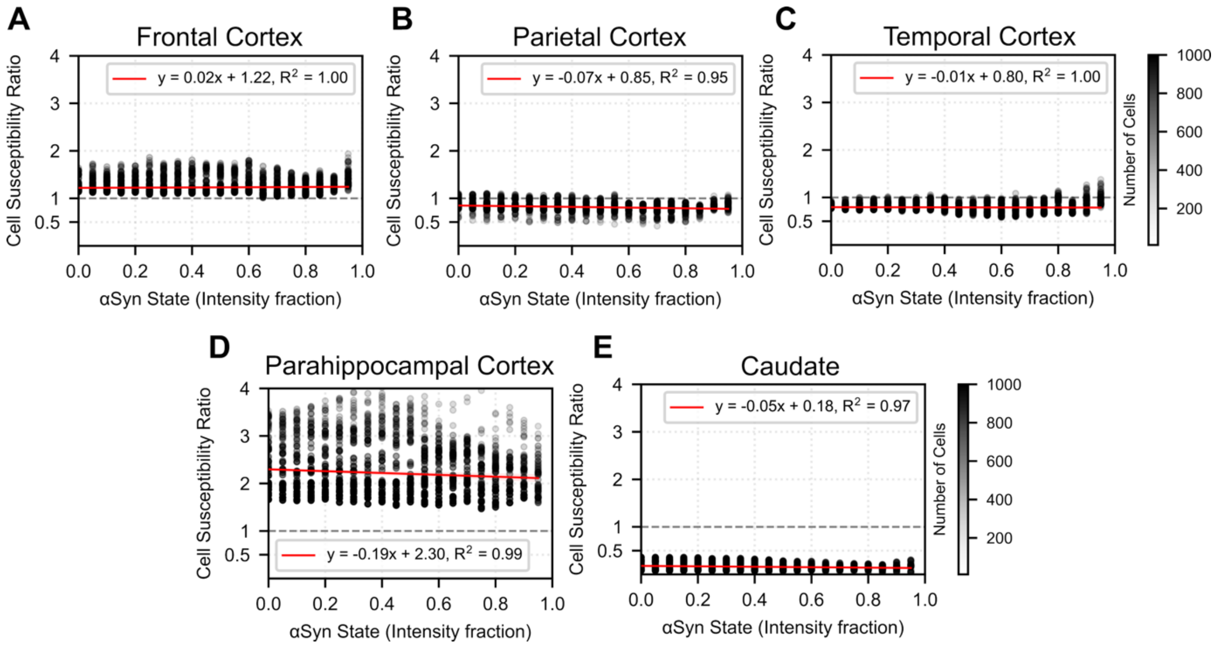


**Figure S8. Cell susceptibility is not dependent on aggregate brightness.** Cell susceptibility ratio (PD/HC) as a function of the intensity fraction of all observed intracellular αSyn aggregates in **A.** frontal cortex **B.** parietal cortex **C.** temporal cortex **D.** parahippocampal cortex and **E.** caudate. Lookup table indicates the number of cells per point on each graph. Multiple points at each value on the x-axis are based on a scanning threshold (See **Section S4**) from T = 0 to T = 3 in 0.05 steps. The y-axis represents the cell susceptibility ratio (See **Figure 3 B**) as the ratio of the fraction of ASCs in PD/ ASCs in HC. Red lines represent simple linear fits weighted by number of cells.

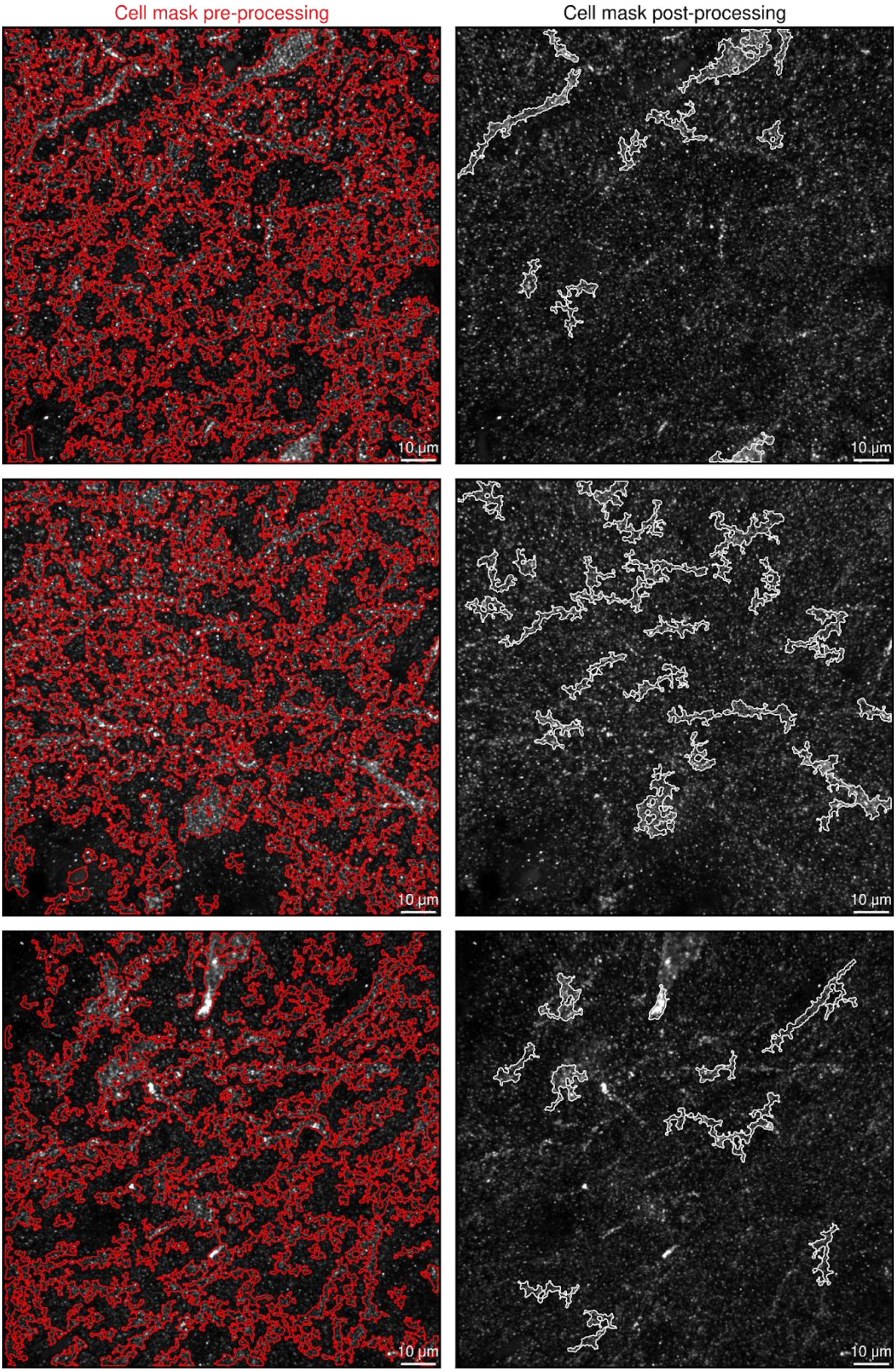


**Figure S9. Cell segmentation is defined by size parameters.** Comparison of un-processed cell mask (left) with processed cell mask (right). Independent unconnected objects of >= 150 and <= 700 µm$^3$ are kept post processing, corresponding to assumed spherical diameters of 6.59–11.02 µm.

### Section S1. Estimation of Lewy Body abundance in Cortical Brain regions

No conclusive quantification of the relative number of Lewy Bodies (LBs) in relation to the number of neurons in neocortical brain regions such as the frontal cortex exists. Few studies look at the fraction of neurons containing LBs, with evidence from the *substantia nigra* showing an average of 3.7% of neurons containing a LB.[7] Across 45 cases, Mattila *et. al.*[8] counted LBs in neocortical regions. On average, they observed 9.22 LBs in the frontal gyrus, 13.8 in the temporal gyrus, 13.58 in the straight gyrus, 7.5 in the precentral gyrus and 6.24 in the angular gyrus, for a total of 50.34 LBs in neocortical brain regions.[8] According to estimates, the approximate number of neurons in the neocortex is 21 billion.[9] Assuming the total number of neurons in the neocortex and the observed number of LBs in neocortical gyri, $2.397 \times 10^{-9}$ % of neurons in the neocortex contain a LB in Parkinson's Disease (*i.e.* 1 in ~400 million neurons). More conservatively, using the total number of LBs observed by Mattila *et. al.*[8] in not just neocortical regions and presuming that all gyri mentioned contain 20% of the total number of neurons in the neocortex, this fraction still is exceedingly small at $79.33/4{,}200{,}000{,}000 = 1.888 \times 10^{-8}$ (*i.e.* 1 in ~50 million neurons). This evidence shows that Lewy Bodies are an exceedingly rare event at the neuronal cell population level. We urge caution in the interpretation of this value; however, it does serve to demonstrate the rarity of LB formation.

### Section S2. Brain region choice rationale after Braak

The brain regions Frontal Cortex (FC), Parahippocampal Cortex (PC) and Caudate Nucleus (CAU) were chosen as representative brain regions for no Lewy pathology (LP), mild LP and moderate LP, respectively, according to Braak.[5,6] In the Braak staging used for neuropathological characterization of the samples, the Caudate Nucleus is moderately affected with substantial evidence of LP due to its direct proximity to the *Substantia Nigra.* The Parahippocampal Cortex first becomes affected in Braak stage 4, meaning LP deposition is currently ongoing in samples from patients classified as Braak 3/4. Neocortical lobes, such as the frontal, parietal and temporal cortices first show sparse LP in Braak stage 5–6, meaning very few to no LBs are expected in Braak 3/4.

### Section S3. Antibody requirements posed by ASA-PD

The previously established ASA-PD protocol for single diffraction-limited αSyn aggregate detection necessitates specific probe characteristics in order to enable the detection of these aggregates.[2] The detection of aggregates can only be possible if local intensity maxima arise as a consequence of probe (here antibody) binding to their targets. If an antibody binds αSyn with a high specificity and affinity, it is likely to abundantly bind monomeric protein which is abundant in brain samples, thereby effectively increasing the overall background signal. A sufficiently low background signal is required in order to accurately detect dim local maxima produced by antibodies binding nanoscale assemblies with approximately >3 antibody binding events per oligomer. If the background produced by an antibody due to its binding of monomeric protein is above the intensity of a nanoscale aggregate, the antibody is not suitable for the ASA-PD protocol, despite its high sensitivity and affinity. Equally, conditions for antibodies are set by the RASP pipeline which is used to detect and quantify technical true positives from the raw microscopy data.[1] Accurate quantification of diffraction-limited local maxima which are oligomers of αSyn necessitates sufficient spatial separation between maxima, or sparsity. Selection of an antibody targeting pS129 αSyn allowed for sub-setting of all possible αSyn epitopes in the sample which increased the visual sparsity of signal compared to an antibody targeting a generic epitope on the protein. This is shown in **Figure S2** which highlights that of the antibodies tested, only anti-pS129 gives the sufficient sparsity necessary for our imaging protocol. However, this evidence does not preclude the use of new probes against αSyn with the abovementioned protocols, providing they meet the necessary conditions.

### Section S4. Summary bar graph computation

The summary bar graphs in **Figure 3 A + Figure 3 B** show the mean ratio of the HC histogram over the PD histogram computed bin-by-bin with error bars are the standard deviation across bins. For Figure **3 B**, this is achieved by computing the histogram of PD/HC ratios of ASCs (%).

### Section S5. Scanning threshold application

**Figure 2 C.** shows the quantification of aggregation susceptible cells (ASCs) through a scanning threshold on the distribution of intracellular αSyn concentrations of neurons for each brain region in PD patients and HCs. Essentially, we aim to test whether there are proportionately more cells containing a higher aggregate concentration in PD than in HCs. Classically, the 1.5 × interquartile range (IQR) rule has been used to define an outlier, *i.e.* an unexpectedly high value, given an approximately normal distribution.[10] However, 1.5 × IQR presumes an approximately normal distribution and is less robust to skewed distributions and lower n-numbers. Therefore, we chose to apply a scanning threshold to the distribution of aggregate concentration per cell values in each brain region. For each brain region, the scanning threshold starts at the mean value of a joint PD + HC distribution of aggregates per neuron. Then, the threshold is increased in increments of $\mu + (0.01 \times \text{IQR})$. At each increment of threshold, the proportion of ASCs is quantified for both the PD and HC distributions separately by dividing the number of cells above the current threshold value of aggregate concentration over all cells observed in the patient group's brain region. This approach ultimately yields a more robust observation of the overall degree of separation between the PD and HC distributions and the relative abundance of high-concentration ASCs as a function of the threshold that can be seen in **Figure 3 B.** Importantly, this data shows that the determination and setting of a threshold to identify an ASC interacts with the relative difference in prevalence of ASCs when comparing the PD and HC groups. As thresholds are set at increasingly extreme values of the distribution ($\mu + {\sim}2.0 \times \text{IQR}$), very few cells have as extreme values in both groups, and the data becomes less reliable and more error prone as sampling is limited. Reliably, $\mu + 1.5 \times \text{IQR}$ shows the most robust difference observed between ASC abundance in PD and HC (**Figure 3 B.**).

### Section S6. Kernel density estimates

In **Fig. 3B** we utilised a kernel density estimation (KDE) plot, specifically the kdeplot function from seaborn.[11] This was utilised as, with the relatively low numbers of data points for a 2D histogram, outliers visually skew the distribution observed whilst contributing very little to the actual form of the distribution. A KDE plot is far less sensitive to these issues, and highlights the underlying shape of the distribution observed. KDEs are in essence the basis of violin plots, and due to their nonparametric nature and reliability in presenting the underlying forms of distributions,[12] we chose to use them here.

### Section S7. Data processing code availability

Code used in this paper is available at (https://doi.org/10.5281/zenodo.16411305). The code package, “pyRASP_copy_for_paper.zip”, contains the python code used in the paper for image analysis and for postprocessing of the image analysis. This postprocessing involves determining if a single oligomer is inside or outside of a cell and determining, for single cells, [αSyn Aggregate]$_{cell}$. A notebook in this zip folder takes the user through the process of loading in raw data and determining [αSyn Aggregate]$_{cell}$. A comprehensive database file of all analysed data is also provided.

| Case ID | Sex | Age of Onset | Age at Death | Disease Duration | PMI | NPD | αSyn Braak | Tau Braak | Aβ Thal |
|---|---|---|---|---|---|---|---|---|---|
| PD1 | F | 65 | 75 | 10 | 14 | LBDBS | 4 | 2 | NA |
| PD2 | F | 77 | 86 | 9 | 22 | LBDBS | 4 | 2 | NA |
| PD3 | M | 66 | 72 | 6 | 9 | LBDBS | 4 | 2 | NA |
| PD4 | F | 71 | 82 | 11 | 16 | LBDBS | 3 | NA | NA |
| PD5 | M | 70 | 85 | 15 | 16 | LBDBS | 4 | 2 | NA |
| PD6 | M | 62 | 78 | 16 | 11 | LBDL | 4 | NA | NA |
| PD7 | M | NA | 86 | 19 | 16 | LBDBS | 3 | 1 | NA |
| PD8 | M | NA | 81 | 17 | 22 | LBDL | 4 | 2 | NA |
| PD9 | F | NA | 76 | 25 | 8 | LBDL | 4 | 1 | 1 |
| PD10 | M | NA | 77 | 1 | 24 | LBDL | 4 | 2 | 3 |
| PD11 | M | NA | 69 | 16 | 13 | LBDBS | 3 | 2 | 2 |
| PD12 | M | NA | 73 | 7 | 24 | LBDBS | 3 | 1 | 3 |
| PD13 | M | NA | 78 | 21 | 19 | LBDL | 4 | 1 | 1 |
| PD14 | M | NA | 91 | 17 | 6 | LBDBS | 4 | 2 | NA |
| HC1 | M | NA | 71 | NA | 29 | Control | NA | NA | NA |
| HC2 | M | NA | 88 | NA | 8 | Control | NA | NA | NA |
| HC3 | F | NA | 92 | NA | 24 | Control | NA | NA | NA |
| HC4 | F | NA | 87 | NA | 12 | Control | NA | NA | NA |
| HC5 | M | NA | 90 | NA | 12 | Control | NA | NA | NA |
| HC6 | M | NA | 87 | NA | 31 | Control | NA | NA | NA |
| HC7 | M | NA | 75 | NA | 24 | Control | NA | NA | NA |
| HC8 | F | NA | 84 | NA | 22 | Control | NA | NA | NA |
| HC9 | M | NA | 75 | NA | 17 | Control | NA | NA | NA |
| HC10 | F | NA | 89 | NA | 13 | Control | NA | NA | NA |
| HC11 | M | NA | 82 | NA | 48 | Control | NA | NA | NA |
| HC12 | M | NA | 75 | NA | 12 | Control | NA | NA | NA |
| HC13 | F | NA | 89 | NA | 20 | Control | NA | NA | NA |

**Table S1.** Case demographics of study cases. Parkinson’s disease (PD) cases and neurologically normal control (HC) cases. PMI – Post-mortem interval; NPD – Neuropathological diagnosis; LBDBS – Lewy Body Disease Brainstem predominant; LBDL – Lewy Body Disease Limbic predominant; NA – Data not available.
